# Integrating long-range connectivity information into de Bruijn graphs

**DOI:** 10.1101/147777

**Authors:** Isaac Turner, Kiran V Garimella, Zamin Iqbal, Gil McVean

**Affiliations:** Wellcome Trust Centre for Human Genetics, Oxford, OX3 7BN, UK; Big Data Institute, Li Ka Shing Centre for Health Information and Discovery, Oxford, OX3 7LF, UK; European Bioinformatics Institute (EMBL-EBI), Wellcome Genome Campus, Hinxton, CB101SD, UK

**Author notes:** These authors contributed equally to the work (Isaac Turner, Kiran V Garimella).

## Abstract

**Motivation:** The de Bruijn graph is a simple and efficient data structure that is used in many areas of sequence analysis including genome assembly, read error correction and variant calling. The data structure has a single parameter *k*, is straightforward to implement and is tractable for large genomes with high sequencing depth. It also enables representation of multiple samples simultaneously to facilitate comparison. However, unlike the string graph, a de Bruijn graph does not retain long range information that is inherent in the read data. For this reason, applications that rely on de Bruijn graphs can produce sub-optimal results given their input.

**Results:** We present a novel assembly graph data structure: the *Linked de Bruijn Graph (LdBG)*. Constructed by adding annotations on top of a de Bruijn graph, it stores long range connectivity information through the graph. We show that with error-free data it is possible to losslessly store and recover sequence from a Linked de Bruijn graph. With assembly simulations we demonstrate that the LdBG data structure outperforms both the de Bruijn graph and the String Graph Assembler (SGA). Finally we apply the LdBG to *Klebsiella pneumoniae* short read data to make large (12 kbp) variant calls, which we validate using PacBio sequencing data, and to characterise the genomic context of drug-resistance genes.

**Availability:** Linked de Bruijn Graphs and associated algorithms are implemented as part of McCortex, available under the MIT license at https://github.com/mcvean/mccortex.

## 1 Introduction

Most efforts to discover genetic variation in populations begin with alignment of high-throughput sequencing (HTS) data to a high-quality reference genome for the organism under study. This approach works well for regions with low divergence from the reference haplotype. However, many biologically interesting loci reside in regions of high divergence. For example, antigenic genes in *Plasmodium falciparum*, *Trypanosoma brucei*, and other pathogens often exhibit non-allelic homologous recombination underlying mechanisms of immune escape (Freitas-Junior *et al.*, 2000; Artzy-Randrup *et al.*, 2012; Jackson *et al.*, 2012). Similarly, structural mutations, such as rearrangements and amplifications, can promote tumourigenesis through dysregulation of oncogenes or down-regulation of tumour suppressors (Difilippantonio *et al.*, 2002; Aguilera and Gómez-González, 2008). More generally, variants may be difficult to identify and characterise when the altered haplotype differs substantially from the reference, and other regions of interest reside in sequence absent from the reference sequence altogether. For example, in 13 isolates of the diploid coccolithophore *Emiliania huxleyi*, 8 to 40 Mbp of the approximately 142 Mbp genome were found to be isolate-specific; up to 25% of genes were found to be absent from the reference sequence (Read *et al.*, 2013). In these scenarios, reads may fail to map to the reference, preventing the analyst from inspecting biologically interesting variation. Alternatively, reads may map incorrectly, misleading the analyst to consider variation where none exists (Ribeiro *et al.*, 2015).

One mitigation of this inadequate reference problem is to augment the reference with known variation and alternative alleles to improve read mapping (Schneeberger *et al.*, 2009; Huang *et al.*, 2013; Weisenfeld *et al.*, 2014; Dilthey *et al.*, 2015). Such approaches commonly convert flat reference genomes into a graph structure, effectively mapping reads to all references simultaneously and choosing the path that best fits the data. In a study mapping to a fragmented human assembly, Limasset *et al.* (2016) found that mapping to a reference graph instead of flat contigs led to a 22% increase in the number of reads that map uniquely.

*De novo* assembly offers a means to overcome some of the limitations of reference-based analyses. Rather than aligning reads to a reference, reads are aligned to one another. These alignments are encoded in a graph data structure, a collection of “vertices” encapsulating sequence data and “edges” representing overlaps of different sequences (Myers, 1995). Graphs from different samples (and any reference) can then be compared to discover variation directly (Bateman *et al.*, 2016). Should the variation be in a locus unrepresented in the reference genome, the graph-based comparison can still capture the event (Iqbal *et al.*, 2012).

The most common sequencers in use today (second-generation) produce tens of millions of short reads (typically 75 to 150 bp in length) per sequencing run (Goodwin *et al.*, 2016). It is common to assemble such data using a so-called “de Bruijn” graph approach (de Bruijn, 1946; Pevzner, 1989; Idury and Waterman, 1995). Vertices are constrained to be fixed-width substrings of length *k* (or “*k*-mers”). Edges represent observed sequence adjacencies in the reads. With sufficient coverage, overlaps are implicitly encoded because two reads which overlap will share *k*-mers. Thus the graph is built up one read at a time at the cost of storing the graph in memory. Graphs of multiple individuals can be compared in memory (Iqbal *et al.*, 2012). However, there is a penalty for this approach: long-range information in the read is sacrificed. This is particularly problematic as genomes tend to have many repetitive regions and without context it is often not possible to determine the origin of a random *k*-mer (Pevzner *et al.*, 2004; Miller *et al.*, 2010). However, as *k* increases, so does the specificity of its location. String graphs address the issue of storing long-range information by avoiding the read fragmentation step and instead find explicit overlaps between reads. Unfortunately string graphs are not well suited to multi-sample comparison and have a high per-sample memory cost (Bonizzoni *et al.*, 2016).

We start by describing the de Bruijn graph and its benefits compared to the string graph. We then describe an augmentation (LdBG) that allows long-range information to be kept. Theoretical results and simulations are used to characterise its properties. We demonstrate its value by application to variant discovery and characterisation of genomic context for drug resistance genes in *Klebsiella pneumoniae*. Finally, we consider the possibility of using such structures for regular analysis of human-scale genomes.

## 2 Background

### 2.1 Definitions and notation

DNA sequences are strings over the alphabet {*A*, *C*, *G*, *T*}. We denote a DNA sequence as *S* = *S*_1_,…,*S_S_* where *S* is the length of the sequence. *S*[*i,j*] is sequence *S_i_*,…,*S_j_*. *S*^′^ is the reverse of *S* (*S_S_*,…,*S*_1_). 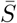 is the reverse complement of S. A *k*-mer is a sequence of length *k* over the alphabet {*A*, *C*, *G*, *T*}.

### 2.2 Assembly graphs

An assembly graph is any graph where the the vertices represent sequence and edges represent overlaps or adjacencies between those sequences. An assembly graph may not have parallel edges (not a multigraph). Traversing a vertex *v* backwards 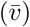 gives the reverse complement of the sequence it represents. *deg*^−^(*v*) is the indegree and *deg*^+^(*v*) is the outdegree of vertex *v*. A path through the graph is a list of adjacent vertices with edges between them. The reverse of path *P* = *v*_1_,…,*v_n_* is 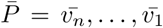. A *unitig U* = *v*_1_,…,*v_n_*, is a maximal path such that *deg*^−^(*v_i_*) = *deg*^+^(*v_i_*) = 1 for 1 < *i* < *n* and *deg*^+^(*v*_1_) = *deg*^−^(*v_n_*) = 1 if *n* > 1. The maximal property means the path cannot be extended without violating these conditions.

### 2.3 De Bruijn Graphs

A de Bruijn Graph *G*(*k*) is an assembly graph, constructed from a set of sequence reads *R* and defined by {*V, E*} where *V* is a set of vertices representing *k*-mers and *E* a set of edges between those *k*-mers. De Bruijn Graphs are constructed by breaking input reads into overlapping *k*-mers that are added to the graph. With one *k*-mer starting at every base, a read of length *r_i_* will give *r_i_* − *k* + 1 *k*-mers. A count is kept of how many times a given *k*-mer was seen in the input reads, called *k*-mer coverage. Edges are added between two *k*-mers if they share an overlap of *k* − 1 bases. Some implementations additionally require that *k*-mers are seen overlapping by *k* − 1 bases in the read data, in order to have an edge between them.

Due to the double stranded nature of DNA and the fact that we don’t know which strand a read originated from, storing all *k*-mers from reads results in *k*-mers occurring separately in the graph in both their forward and reverse complement orientations. To overcome this it is common to store only the lexically lower of each *k*-mer *X* and its reverse-complement 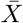 (Zerbino, 2010). Requiring that *k* is odd prevents a *k*-mer from being its own reverse complement (a DNA palindrome). When visiting a vertex in the de Bruijn graph we can visit it in its forward or reverse complement orientation. The orientation in which we arrive at it determines if we leave by its out- or in- edges (forward, reverse respectively).

A de Bruijn graph only stores connectivity information one base either side of a given *k*-mer. This means that for three adjacent *k*-mers in the graph (*v_a_, v_b_, v_c_*), there is no information about how the first and third are connected if the middle *k*-mer has *deg*^−^(*v_b_*) > 1 and *deg*^+^(*v_b_*) > 1. This graph motif is known as a ‘tangle’ and is caused by the graph collapsing down at a repeat and splitting out again afterwards. De Bruijn graphs collapse down at repeats in the genome of lengths ≥ *k*. It is not possible to traverse a dBG past a tangle, even if the input reads are long enough to resolve it (i.e. pair-up *k*-mers going-into and coming-out of it). This makes analyses that use a de Bruijn graph sensitive to the parameter *k*.

While increasing *k* can overcome the problem of short repeats, it also has the effect of reducing the number of *k*-mers given by each read and increases the number of *k*-mers lost to each sequencing error. Both these effects reduce *k*-mer coverage, which is determined by the *k*-mer size, the read length and the error rate (Iqbal *et al.*, 2012). As *k*-mer coverage drops, read overlaps are lost and gaps in coverage increase. Together with tangles, coverage gaps interrupt assembly and shorten contigs.

Picking a value for *k* is ultimately a trade-off. It is common to run analyses multiple times with different values of *k* and pick the best results according to a quality metric (e.g. assembly N50 or number of variants called) (Iqbal *et al.*, 2013). Alternatively the genome and read data can be sampled to estimate which value would be optimal (Zerbino, 2010).

The dBG can be augmented to support multiple data sets, providing a single data structure to describe and compare the genomes of many individuals (Iqbal *et al.*, 2012). Graphs are built separately for each data set *c* ∊ *C* and merged post-construction. The merge produces a union graph *G_u_* = {*V_u_, E_u_*}, where 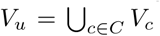 and *E_u_* = {*E_c_* : *c* ∊ *C*}. Each *k*-mer stores which samples it was seen in. We refer to *c* as *colour*, a generic term that can mean a distinct individual, pooled population or a specific data set on a single individual (e.g. tumour/normal), depending on analysis context. We shall refer to this structure as a *multi-colour de Bruijn graph*.

De Bruijn graphs are used in many areas of sequence analysis, including in mapping-based calling, as in the local alignment step of the variant caller Platypus (Rimmer *et al.*, 2014), in de novo assembly as in Velvet (Zerbino and Birney, 2008) and ABySS (Simpson *et al.*, 2009), and in de novo assembly for variant calling (Iqbal *et al.*, 2012).

Recently there has been work on implementing low memory dBG construction (Chikhi *et al.*, 2014) and representations (Conway and Bromage, 2011; Rizk *et al.*, 2013; Chikhi and Rizk, 2013; Bowe *et al.*, 2012). These have both provided great improvements over the naive hash table based implementation, extending the contexts in which dBGs can be used.

### 2.4 String graphs

A String graph is an assembly graph where the vertices represent the input reads and the edges are maximal non-transitive overlaps between them (Myers, 2005). The set of reads is reduced to remove reads contained within other reads. A naive String Graph implementation would take *O*(*N*^2^) time to compare all pairs of reads to find overlaps, before removing contained reads and transitive edges. Simpson and Durbin (2010) showed that it is possible to construct a string graph in linear time, by first generating an FM-index of the input reads *R* and an FM-index of their reverse *R*′. Alternatively a single index can be constructed containing *R* and 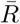 (Li, 2012).

The FM-index (Ferragina and Manzini, 2000) is a data structure for compression and fast string searching. The FM-index of a set of strings *S* facilitates searching for a query *Q* in time *O*(|*Q*|). Construction of the index has time and memory complexity *O*(|*S*|). The final index has roughly the same size *S*, but can be efficiently compressed with run-length encoding.

FM-index construction has high memory requirements for high coverage data sets as all reads must be loaded into memory. This requires building indices for subsets of the reads and recursively merging the indices (Li, 2014a). In comparison, dBG memory is mostly determined by genome size, with only the errors in high coverage data causing a slower increase in memory requirements. Similar memory efficiency advantages are seen when building multi-colour dBGs from the same species.

Since it is constructed from the reads without breaking them up, a string graph is an assembly graph that stores all the connectivity information contained in the single-ended input reads (Myers, 2005). String graphs do not naturally lend themselves to storing information on read pairs, although one such data structure has been proposed (Chikhi and Lavenier, 2011).

### 2.5 The linked de Bruijn graph

We propose a new assembly graph data structure called the Linked de Bruijn Graph (LdBG). Defined as *LG*(*k*) = (*V*, *E*, *L*) where *V*, *E* are defined as in a de Bruijn graph. *L*(*v*) is a set of paths through the graph that start at vertex *v* ∊ *V*. We call these paths *links*. Each of these links *l* ∊ *L*(*v*) is stored as a list of junction choices that when followed, starting from vertex *v*, recreate the path. Graph traversal is the same as with a de Bruijn graph, with the extension that when we visit a vertex *v*, we pick up the links associated with it: *L*(*v*). The links held during traversal record how many edges ago they were picked up, a value we call link “age”. Only when we reach a bifurcation in the graph do we consult the links currently held. We follow the next junction choice of the oldest link as this provides the most context as to where we are in the genome. Should we have more than one oldest link and they disagree, we halt traversal. An illustration of links resolving a cycle is shown in Figure 1.

**Figure 1:**
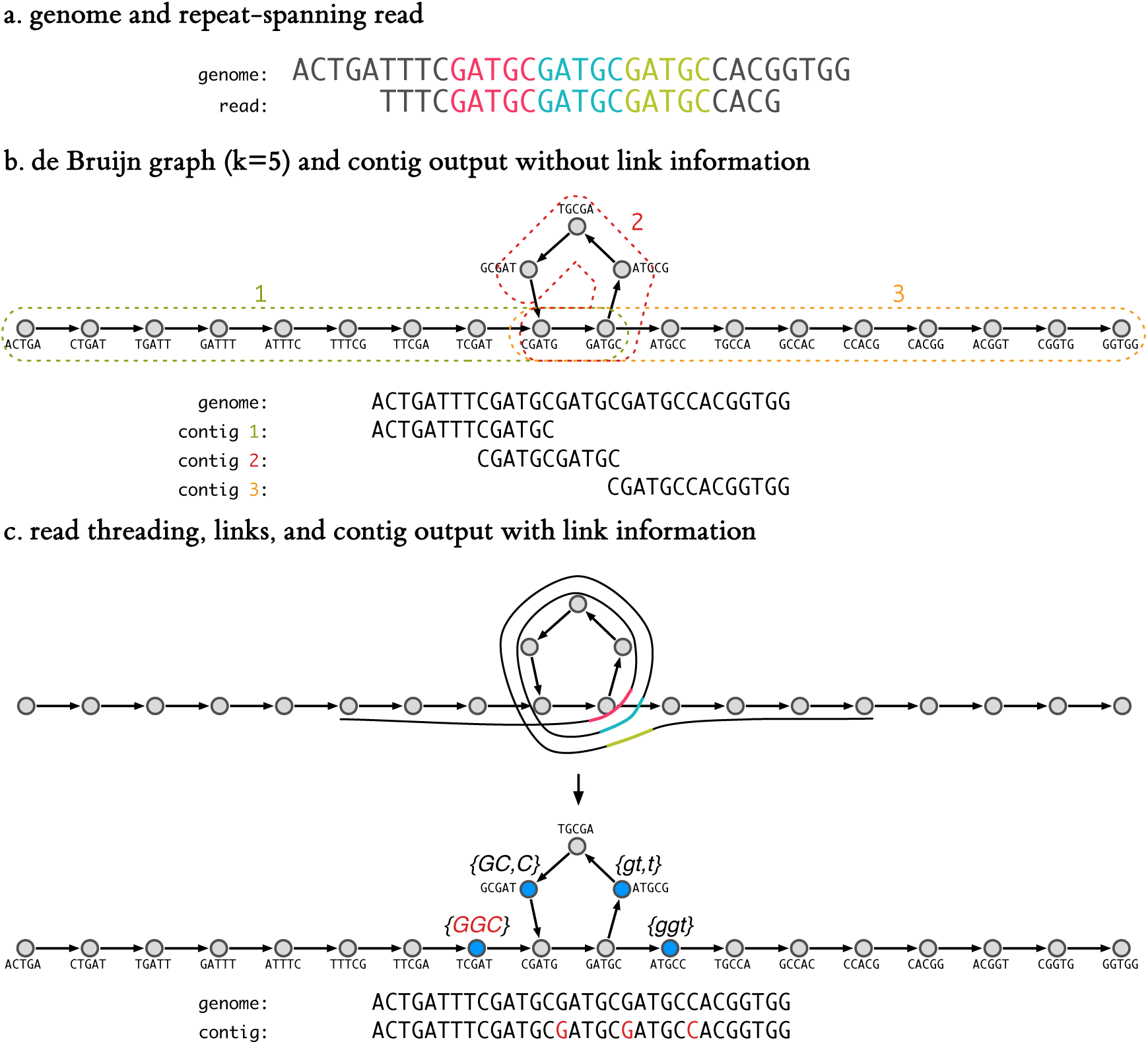
Utility of link information in traversing a graph cycle. (a) A 32-bp genome and a 23-bp read, each containing three (colour-coded) repeats of the 5-mer, GATGC. (b) The resulting de Bruijn graph (*k* = 5) with a repeat cycle, constructed from the genome sequence. The *k*-mers grouped by dashed boxes indicate the result of graph traversals to emit contigs, with final sequences written below and positioned along the input genome for clarity. (c) Reads are “threaded” (aligned) through the graph (top); the repeated *k*-mers are colour-coded. The alignment information is distilled to a set of junction choices to make when navigating the graph and stored as annotations on *k*-mers preceding junctions (middle). Multiple links are separated by a comma. Uppercase (lowercase) links indicate the choices to be made when traversing forwards (backwards). A *k*-mer’s links are picked up when we visit it. When we reach a junction, the next edge suggested by the oldest link(s) is taken, links that disagree are dropped, all remaining links trim off a junction choice and exhausted links are also dropped. The resultant contig recapitulating the entire genome is shown (bottom). Highlighted bases indicate the junction choices originating from the left-most link.

As with a de Bruijn graphs we can look up any *k*-mer or edge between *k*-mers in time *O*(1) and we can start graph traversal from any *k*-mer. As in a multi-coloured dBG, a multi-coloured LdBG stores which samples have which *k*-mers and links.

A LdBG is a lossless representation of a genome when generated from error-free reads, as long as the genome starts and ends with unique *k*-mers, there are no *k*-mer coverage gaps and each repeat is spanned by at least one read (proof in online supplement). This is true regardless of the value of *k*.

In constructing a LdBG we are effectively compressing reads against the de Bruijn graph. However, since read start/end positions are not important for assembly we do not store them, so although it is possible to recover the underlying genome (losslessly) through assembly, it is not possible to recover the original set of input reads.

Reads used to annotate the graph do not need to have been used to construct the de Bruijn graph. Sets of links may be merged by loading them together at runtime. We give an example of the utility of such a construction in the applications section below.

## 3 Methods

### 3.1 de Bruijn Graph construction

Each input read *r* is broken into *r* − *k* + 1 overlapping *k*-mers (*v*_1_,…, *v_n_*) which are added to the graph. If a *k*-mer already exists in the graph, we increment its coverage. Edges are added between *v_i_* and *v_i_*_+1_ for all 1 ≤ *i* < *n*.

To remove *k*-mers due to sequencing error, unitigs with median *k*-mer coverage below *T* are removed, where *T* is a user-specified threshold. If not specified, a threshold *T* is picked such that the expectation of a *k*-mer with coverage *T* being an error is < 10^−3^ (see online supplement).

Graph tips, that is unitigs (*v_i_,*…,*v_j_*) with *deg*^−^(*v_i_*)+*deg*^+^(*v_j_*) < 2, are the result of sequencing errors near the end of reads and gaps in coverage. Tips are removed if they are shorter than a user specified value, the default being *k*, the maximum number of erroneous *k*-mers generated by a single-base sequencing error near the end of a read.

### 3.2 Read-to-graph alignment

Reads are aligned to the de Bruijn graph one-at-a-time and in doing so are error-corrected. For a read *r*, we look up each of its *k*-mers, resulting in a list of *k*-mers that describe a path through the graph. There may be gaps in this path due to *k*-mers removed from the graph during *k*-mer error cleaning (or if the read was not used in dBG construction). Gaps are closed by walking the graph between the *k*-mers either side of the gap (*v_i_* and *v_j_*). If we cannot traverse from *v_i_* to *v_j_*, we attempt going from *v_j_* to *v_i_*. Should such a traversal succeed (giving *k*-mers *v*_1_,…,*v_k_*) and *k* ~ *j* − *i*, the *k*-mers *v*_1_,…,*v_k_* are used to fill in the gap in the path between *v_i_* and *v_j_*. This error step is sequence agnostic in that it does not compare the new *k*-mers (*v*_1_,…,*v_k_*) to the read *k*-mers it is replacing (*v_i_*_+1_, …,*v_j_*_−1_). This speeds up the error correction step and ensures it does not make assumptions about the error process of the input sequence data. The output of the alignment step is a set of sequences that perfectly match the de Bruijn graph *k*-mers i.e. they describe a path through the graph.

Gaps between paired-end reads are treated like gaps in reads caused by sequencing errors. LdBG naturally captures information from paired end reads once the insert gap is filled. Links can be generated in two passes: first with single-end reads against a dBG to create a LdBG; then with paired-end reads against the LdBG. This allows the single-ended read links to be used to aid traversal between read pairs.

### 3.3 Link annotation

A link is a path, starting from a given *k*-mer and stored as a series of junction choices. The function *J*(*v_i_*,…,*v_j_*) takes a path and returns the junction choices it describes.

Given a path *P* = *v*_1_,…,*v_n_* through the graph, we identify the maximum *j* such that *deg*^+^(*v_j_*) > 1; 1 < *j* < *n*. Then for each *i* such that *deg*^−^(*v_i_*) > 1; 1 *< i* ≤ *j*, we add a link to vertex *v_i_*_−1_: *L*(*v_i_*_−1_) ← *L*(*v_i_*_−1_) ∪ *J* (*v_i_*_−1_,…,*v_j_*_+1_). Link annotation is repeated for the reverse path *P*. Link counts record how many times a given link is seen in a sample starting at a particular *k*-mer.

Links are cleaned by building a tree of links *L*(*v*), and trimming junction choices with coverage below threshold T. This link cleaning threshold is determined by applying the same model as used for *k*-mer cleaning to the link coverage distribution of the first junction choice of all links.

### 3.4 Implementation

We have implemented the LdBG data structure and associated algorithms as part of `McCortex`, a modular set of multi-threaded programs for manipulating assembly graphs written in C. McCortex supports FASTA, FASTQ, SAM, BAM & CRAM file formats and is released under the MIT license. McCortex has been used as the backend for sequence analysis by Bradley *et al.* (2015).

### 3.5 Multi-coloured linked de Bruijn graphs

Multi-colour LdBGs can be constructed by building single sample LdBGs and loading them together into McCortex. For graph traversal tasks, such as assembly, we only store a single bit per sample per *k*-mer and per sample per link to record which *k*-mers/links are present in each sample. These are stored in a packed bitset. Graph traversal of a colour through a multi-coloured LdBG proceeds as per for a single-sample LdBG, only using links and *k*-mers of the given colour. At coverage gaps, traversal can fall back to using any *k*-mers in the graph (but not other colour’s links).

## 4 Results: simulations

### 4.1 Equivalence of LdBG and input string

To test the lossless recovery of a genome from the LdBG we generated a random 10 kbp haploid genome, ensuring it started and ended with unique 7-mers. We identified the length of longest repeat (*LR*) in our genome. We generated perfect error-free coverage of the genome with a read length of *LR* + 2 starting at each base. We then built a LdBG (*k* = 7) from the reads, assembled contigs and removed contained contigs (those that were substrings of other contigs). After checking that we were left with a single contig, we compared it for an exact match to the original genome. This simulation was run 100 times without fail. With *k* = 7, there are only 4^7^ = 16384 possible *k*-mers, so a random 10 kbp genome will have many repeats that could not be traversed by the unannotated de Bruijn graph.

### 4.2 Correcting errors in reads

To assess the accuracy of our error correction step when aligning reads to the graph, we simulated a haploid 1 Mbp genome (from human GRCh37 chr22:28,000,000-28,999,999). Single-ended 250 bp reads with 50*X* coverage were simulated with a 0.49% empirically distributed sequencing error (reads paired with real MiSeq data, FASTQ scores used as per base error rate). We built a dBG (*k* = 31) and removed tips and unitigs with coverage *<* 7 (automatically chosen). Once reads were aligned to the graph we wrote them to disk instead of generating links. The input reads had 247, 075 (0.49%) errors, the output had 30, 148 (0.06%) errors. Of the bases changed by the error correction step, 99.19% changes were correct.

### 4.3 Sensitivity to word length

Lowering the value of *k* in a dBG raises *k*-mer coverage and reduces coverage gaps but it also reduces the length of the longest repeats that can be traversed. If we improve the ability to resolve repeats with *links*, we hypothesised that we should reduce the assembly performance’s sensitivity to the parameter *k*. Therefore we simulated an assembly task with different *k* values.

We simulated three haploid sequencing data sets from 1 Mbp of human (chr22:28,000,000-28,999,999) using 100 bp single ended reads, each giving 100*X* coverage. First, we generated ‘perfect coverage’ – an error-free read starting at every base. Second, we generated ‘stochastic coverage’ – read starts distributed uniformly across the 1 Mbp genome. Third, we generated ‘reads with error’ – stochastically sampled reads with a uniform 0.5% rate of single base errors.

We assembled these three data sets using a dBG and LdBG at *k* = 21, 31,…, 91. To compare assemblies we used the NG50 metric, defined as the contig length *C* such that contigs longer than *C* sum to at least half of the genome size. NG50 and assembly errors were counted by aligning the contigs to the truth sequence. Single base mismatches were allowed as long as they were flanked by 21 bases that match the graph. Breaks between two alignments were counted as misassemblies. The lengths of aligned sequences were used to calculate NG50.

The NG50 comparisons are shown in Figure 2. In the *“perfect”* data sets reconstructed without links, NG50 rises as *k*-mer size increases. This is to be expected as a longer *k*-mer size essentially encodes more connectivity information. Links, however, encode all available connectivity information at any value of *k*. Thus the linked NG50 value (solid green line) is equal to the best unlinked NG50 (dashed green line) over all values of *k*. The *“stochastic”* data sets (orange) follow a similar pattern, with the exception that the top value of *k* = 91 does not necessarily yield better NG50. As read starts are not available at every single base, some read overlaps are not present and the resulting contig is thus truncated. Finally, the *“error”* data set (blue) shows improved NG50 results when link information is used. When faced with sequencing error, our algorithms are not as readily capable of delivering *k*−independent reconstructions, although using links does improve performance at all values of *k*.

**Figure 2:**
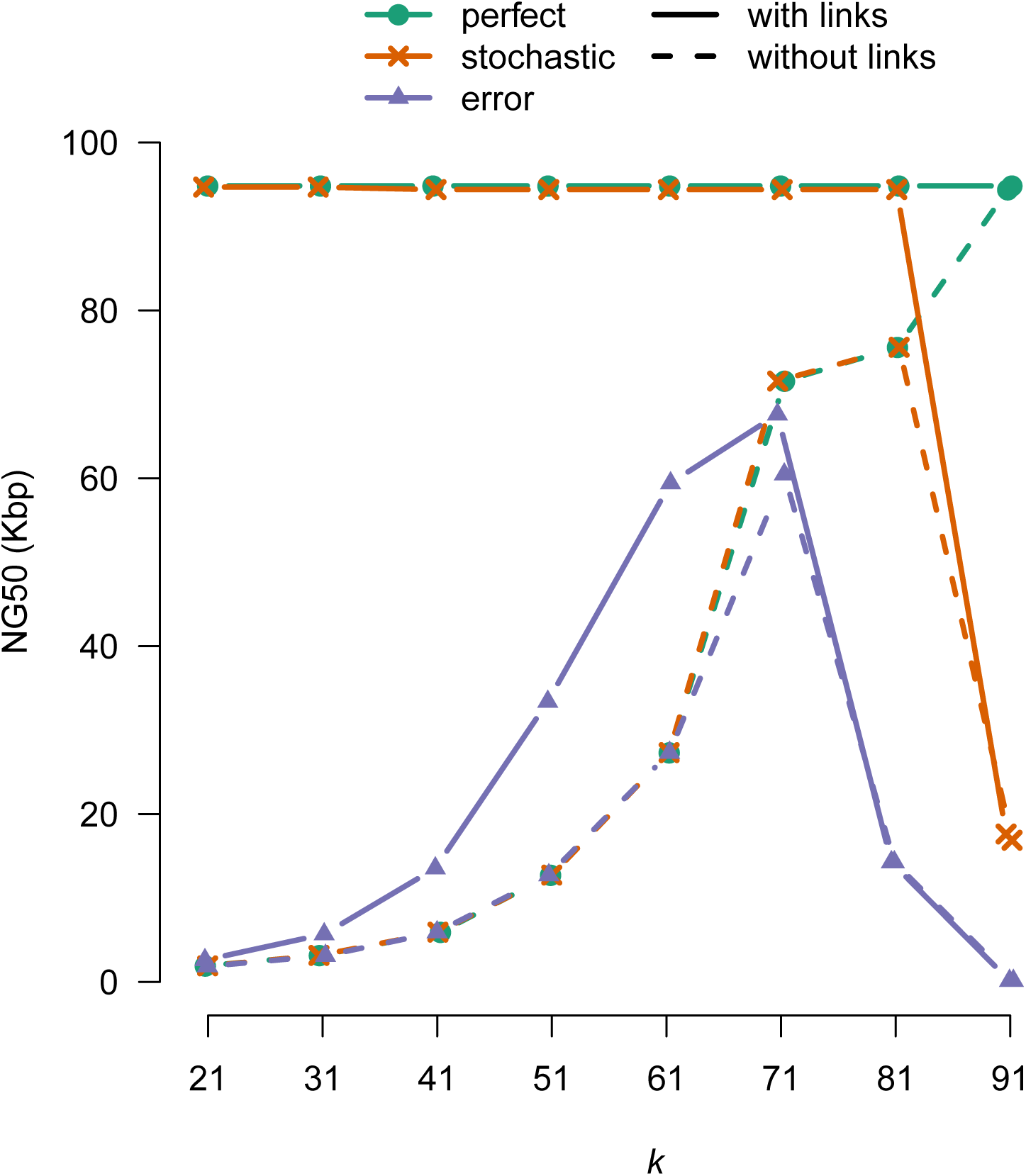
Assembly length metric NG50 on raw de Bruijn graphs (i.e. without links, dashed lines) and linked de Bruijn graphs (i.e. with links, solid lines), as a function of *k*-mer size. Assembling 1 Mbp of sequence (human GRCh37 chr22:28,000,000-28,999,999) with three simulated 100X read data sets: error free 100 bp reads, one read starting at every base (*“perfect”, green*); error free stochastic coverage (uniformly distributed read starts) (*“stochastic”*, orange); an error rate of 0.5% and stochastic coverage (*“error”*, purple).

To explain this behaviour, we note that at low *k*, sequencing errors introduce false edges between true *k*-mers. Since error correction on dBGs use *k*-mer counts rather than edge counts, these false edges do not get cleaned off. We estimated the number of false edges induced at various *k* to be 604, 139, 30, 7, 2, 1, 0, 0, 0 for *k* = 21, 31,…, 91. Each false edge introduces a new bifurcation that may halt traversal. dBG implementations that use counts on edges instead of *k*-mers (as described in Conway and Bromage (2011)) may overcome this issue.

### 4.4 Comparison to string graph assembly

Next, we compared assembly performance between our LdBG implementation and the String Graph Assembler (`SGA`) (Simpson and Durbin, 2010). As stated above, string graphs are notable as their construction is based on the direct computation of read-to-read overlaps, facilitated by an FM-index on the reads. Similar to our work, `SGA` is able to use the full length of the read during assembly and should thus be able to assemble repeats shorter than a read length. However, `SGA` differs from our work in some key ways. Overlaps between reads need not be perfect, but rather are parametrised at run time to accept a minimum overlap of *τ_min_* bases and maximum error rate *ϵ_max_*. Additionally, SGA attempts to correct sequencing errors, rather than discarding them completely. In practice, these two factors may yield longer contigs, but at the expense of accuracy (Bradnam *et al.*, 2013).

The previous simulated haploid genome *“error”* read data set was used. As shown in Figure 2, `McCortex` is sensitive to sequencing error, so we ran `bfc` error correction (Li, 2015) on the reads before assembling. bfc error correction did not have a beneficial effect on `SGA` and was therefore not used. SGA was used to assemble at various values of *τ_min_*. While *τ_min_* and *k* are not identical parameters (without setting *ϵ_max_* = 0) they are still informative in terms of how many bases between reads are required to produce an overlap. Other `SGA` parameters were left to software defaults (most notable for `SGA’s` error correction step, which learns the appropriate *k*-mer coverage cutoff threshold automatically). We did not carry out scaffolding or use paired end information in link construction.

`SGA` and `McCortex` assemblies were compared using the same NG50 and assembly error metrics as above (see Figure 3). When compared to the the true genome, `McCortex’s` NG50 is roughly the same as SGA at low and high *k*, but is much better at *k* = 41,…,71. Assembly errors are about an order of magnitude lower.

**Figure 3:**
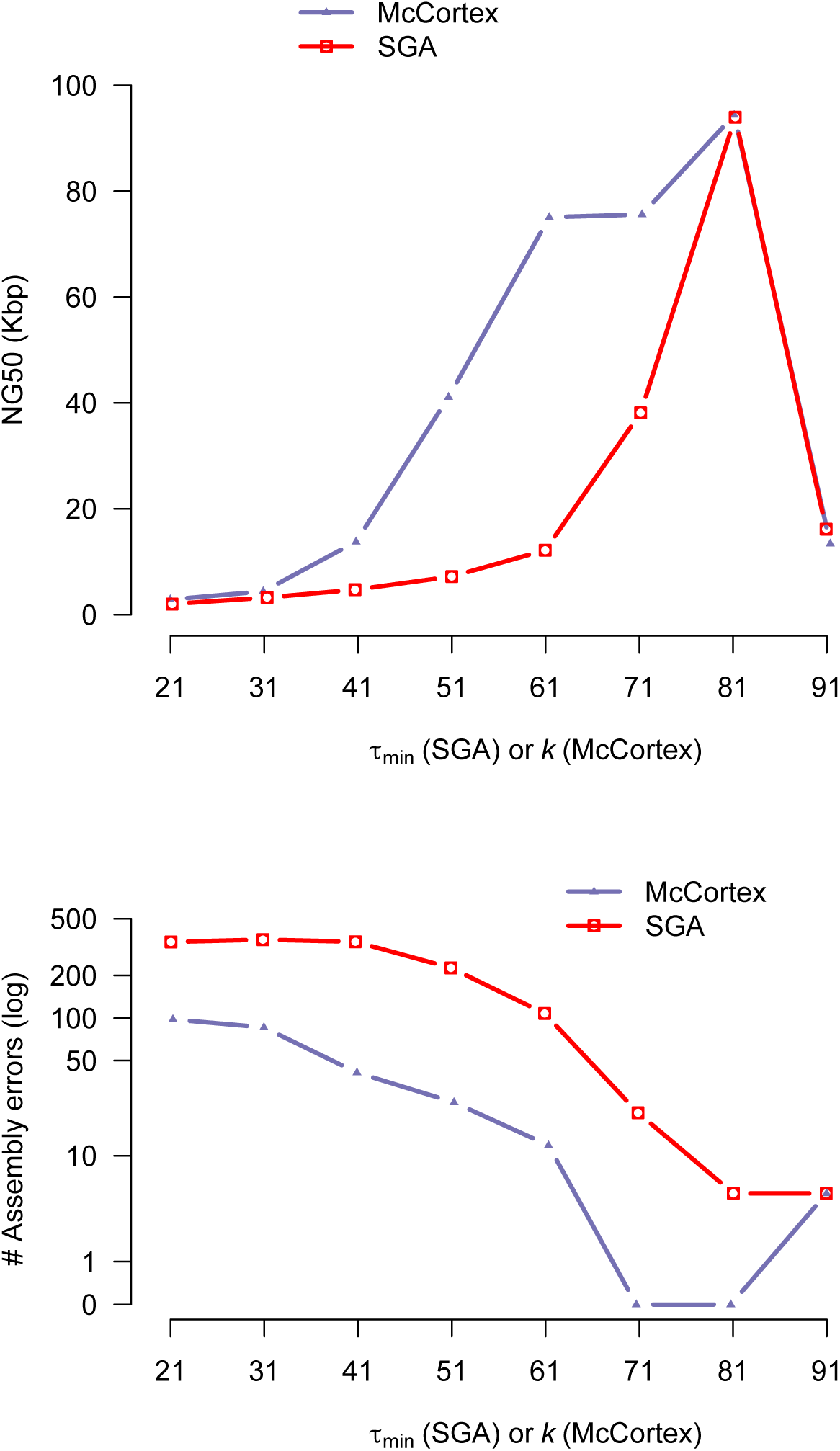
Assembly results for SGA and `McCortex` compared using a simulation on 1 Mbp of sequence (human GRCh37 chr22:28,000,000-28,999,999), using 100 bp reads with a per base error rate of 0.5% and stochastic 100X coverage (“*error*” data set from Figure 2). Top panel: contig NG50 at different parameters of *τ_min_* (SGA) and *k* (LdBG). Bottom panel: number of assembly errors in contigs compared to truth.

## 5 Results: applications

To assess LdBG on real data, we examined short read data from *Klebsiella pneumoniae*, a gram-negative bacteria that usually lives harmlessly in the mouth and gut of humans. However in the event of a weakened immune system, it can establish pathogenic colonies in the lung leading to inflammation and bleeding. It is also found in some cases of urinary tract infections. Antibiotic resistant strains of *K. pneumoniae* have been found in patients. We used McCortex for two tasks where long-range information is likely to be beneficial - finding large differences from a reference and analysis of genomic context for drug resistance genes, which we validated using a PacBio reference assembled for the sample (Sheppard *et al.*, 2016).

### 5.1 Large-variant discovery

As links should provide useful guidance to navigating junctions in a graph, we examined their utility in calling large variants (insertions or deletions greater than 100 bp in length). We implemented a “bubble caller” (named for the characteristic motif produced by a biallelic mutation in a graph wherein paths diverge from one *k*-mer and rejoin at another) and tested it by calling variants in CAV1016, a *K. pneumoniae* isolate for which a high-quality PacBio sequence was available for validation. We constructed dBGs of the canonical reference sequence (GCF 000016305.1 ASM1630v1) and Illumina data for CAV1016. From these, we built LdBGs using the single-end Illumina reads for link construction. We applied our bubble caller to the dBG and LdBGs, allowing for a minimum event size of 100 bp and maximum of 200 kbp, and removing duplicate events. We validated called alleles by aligning the reference and alternate alleles to the canonical reference sequence and CAV1016 PacBio sequence, respectively. The resulting callsets without and with link information are presented in Supplementary Table 1.

**Table 1:** Comparison of KPC-containing contigs to validation data, inferred with and without links

| Isolate | Known plasmid and KPC allele | without *link information* |  |  | with *link information* |  |  |
| --- | --- | --- | --- | --- | --- | --- | --- |
|  |  | Length (bp) | Matches uniquely and correctly | Mismatches, gaps | Length (bp) | Matches uniquely and correctly | Mismatches, gaps |
| CAV1016 | 1, KPC-2 | 3893 | non-unique | 0, 0 | 43630 | yes | 0, 0 |
| CAV1017 | 1, KPC-2 | 3895 | non-unique | 0, 0 | 43628 | yes | 0, 0 |
| CAV1042 | 1, KPC-2 | 2557 | non-unique | 0, 0 | 43669 | yes | 0, 0 |
| CAV1077 | 1, KPC-2 | 3708 | non-unique | 0, 1 (188 *bpdel*) | 15796 | yes | 0, 1 (188 *bpdel*) |
| CAV1142 | 1, KPC-2 | 3892 | non-unique | 0, 0 | 43612 | yes | 2, 0 |
| CAV1145 | 1, KPC-2 | 5158 | yes | 0, 0 | 43643 | yes | 1, 0 |
| CAV1182 | 1, KPC-2 | 2558 | non-unique | 0, 0 | 9722 | yes | 1, 0 |
| CAV1203 | 1, KPC-2 | 3911 | non-unique | 0, 0 | 43670 | yes | 0, 0 |
| CAV1205 | 2, KPC-2 | 5978 | yes | 0, 0 | 14019 | yes | 2, 1 (2 *bpdel*) |
| CAV1207 | 2, KPC-2 | 13249 | yes | 0, 0 | 13991 | yes | 3, 1 (2 *bpdel*) |
| CAV1237 | 1, KPC-2 | 20444 | yes | 1, 0 | 43613 | yes | 2, 1 (8 *bpdel*) |
| CAV1290 | 1, KPC-2 | 5099 | yes | 0, 0 | 29340 | yes | 4, 1 (2 *bpdel*) |
| CAV1292 | 1, KPC-2 | 5094 | yes | 0, 0 | 29337 | yes | 1, 1 (2 *bpdel*) |
| CAV1338 | 2, KPC-2 | 5963 | yes | 0, 0 | 14003 | yes | 2, 1 (2 *bpdel*) |
| CAV1344 | 1, KPC-2 | 4839 | yes | 0, 0 | 9694 | yes | 1, 0 |
| CAV1351 | 1, KPC-3 | 3898 | incorrect | 0, 0 | 43616 | yes | 1, 1 (5 *bpdel*) |
| CAV1360 | 1, KPC-2 | 734 | non-unique | 0, 0 | 43621 | yes | 1, 0 |
| CAV1391 | 1, KPC-2 | 5106 | yes | 0, 0 | 9726 | yes | 1, 0 |
| CAV1576 | 2, KPC-2 | 9161 | yes | 0, 0 | 14003 | yes | 2, 1 (2 *bpdel*) |
| CAV1578 | 2, KPC-2 | 5243 | yes | 0, 0 | 8956 | yes | 0, 0 |
| CAV1597 | 2, KPC-2 | 12627 | yes | 0, 0 | 14009 | yes | 2, 1 (2 *bpdel*) |

Our bubble caller discovered 55 large indels in the dBG and 59 indels in the LdBG. All 55 variants from the dBG callset were recovered in the LdBG callset. The four remaining variants exclusive to the LdBG callset are all insertions of varying size (134, 246, 7, 952 and 11, 946 bp).

That the LdBG-exclusive events should all be insertions (particularly large ones) is perhaps not surprising; in a graphical framework, calling insertions against a high-quality reference sequence with comparatively lower quality Illumina data is expected to be more difficult than calling deletions. With insertions, sequencing error in the study sample will produce spurious paths in the graph, not all of which can be removed successfully, and thus graph traversal from the 5’ to the 3’ end of the alternate allele has many opportunities to fail. With deletions, the graph navigation burden is on the reference allele which should have substantially fewer errors (and thus fewer spurious paths) to confound traversal. Link-informed traversal helps alleviate this insertion/deletion detection bias, enabling the recovery of large events like the 8 and 12 kb events listed above. This improves our access to large variants underrepresented in current variant call sets (Weisenfeld *et al.*, 2014; Li, 2014b).

### 5.2 Reference-link guided assembly

Finally, we show that with links, we can use a panel of reference contigs derived from multiple sources to improve drug resistance locus characterisation in *K. pneumoniae* isolates. As the underlying graphs are considered immutable after construction, links derived from this panel cannot add *k*-mers to a sample. We hypothesised that the links panel could still provide valuable connectivity information where they were consistent with the graph without misleading the assembler in regions where they were divergent. We selected 21 *K. pneumoniae* isolates with known drug resistance status and that carry combinations of two alleles and two plasmid backgrounds at the *K. pneumoniae* carbapenemase (KPC) resistance locus, see Table 1. As references, we constructed links from a panel of four plasmid backgrounds carrying three different KPC alleles: PacBio sequences from two of the 21 isolates (carrying allele KPC-2), a KPC-harbouring plasmid from *E. coli* (carrying allele KPC-3), and a fourth *K. pneumoniae* plasmid known to harbour a resistance allele and background absent from the 21 isolates (carrying allele KPC-5). All accessions are described in Supplementary Table 2. Two assemblies were generated per isolate: one without links, and one with links.

Contigs harbouring the KPC sequence within the 21 isolates were identified by aligning to the KPC-2 allele sequence with LASTZ (Harris, 2007) and extracting the longest such contig from each assembly. These were aligned back to both the reference data sources and the validation data (Mathers *et al.*, 2015). For alignments that ran off the end of a sequence owing to the circular nature of the plasmids, we attempted to shift the contig sequence such that a linear alignment of maximum length was achieved; where this was not possible we have reported the length of the aligned region. The contig selected from each assembly was evaluated for correct KPC allele recovery, correct identification of plasmid background (i.e. sequence context of KPC allele), and mismatches/gaps to the relevant reference sequence. These results are shown in Table 1.

Without link information, we find that in 57% of cases the plasmid background on which the KPC allele resides cannot be identified. In such cases, LASTZ reports alignments of the short contigs with 100% sequence identity to plasmids 1 and 2. Moreover, for the CAV1360 isolate, the aligner determines the background incorrectly as the *E. coli* plasmid due to the presence of KPC-3.

Reconstruction with the link panel provides an order of magnitude increase in contig length over the link-uninformed reconstructions and the inferred plasmid membership matches the Mathers *et al.* (2015) determination in all 21 cases. Reconstructions from two isolates stand out. CAV1351 was known to carry the KPC-3 allele, while all other isolates carried the KPC-2 allele. The link-uninformed assembly produces a contig that maps to the *E. coli* KPC-3 sequence perfectly, but infers the wrong plasmid membership. The link-informed reconstruction, however, produces both the correct plasmid membership and correct allele. In another case, Mathers *et al.* (2015) reported CAV1077 to possess plasmid 1, but with an unspecified sequence alteration. Our reconstruction is able to establish both the correct plasmid membership and identify a 188 bp deletion in the intergenic region upstream from the transposase and downstream from the KPC genes (detected both with and without links). Combined, these analyses demonstrate how using external data sources as a means to guide assembly through “reference links” can lead to highly accurate reconstruction of even complex regions of the genome.

### 5.3 Scaling to human genomes

Finally, to assess the scalability of LdBG to large genomes, we constructed an LdBG (*k* = 31) from paired-end PCR-free human (NA12878) whole-genome Illumina data (2x150bp, 50X, accession ERR194147). Construction has a peak memory usage of 400 GiB; loading the cleaned LdBG into memory requires 70 GiB of RAM (50 GiB for the dBG, 20 GiB for links). Although the memory footprint of construction is high, there is large scope for improvement using compressed and/or disk-based methods (Chikhi and Rizk, 2013).

## 6 Discussion

We have presented a de novo assembly method that addresses the most important limitation of de Bruijn graphs: the ability to leverage long-range connectivity information inherent in read data. While cutting reads into small *k*-mers has long been a useful way of simultaneously computing read-to-read overlaps and overcoming high rates of sequencing error, increasing sequencing quality and read lengths have rendered de Bruijn assembly methods less attractive. String graphs have been successful in incorporating long-range data into assemblies, but sacrifice desirable computational properties of de Bruijn graphs. Our solution, Linked de Bruijn Graphs, combine the connectivity properties of string graphs with the rapid lookup of specific (multi-coloured) *k*-mers. Due to the wide range of uses of dBGs in sequence analysis, we believe this offers a potential improvement to many existing algorithms. Path encoding of reads has been suggested for read compression before (Conway and Bromage, 2011; Kingsford and Patro, 2015). However we believe this is the first implementation to use it for multi-colour assembly that can scale up to large mammalian genomes on modern computer hardware.

We have shown that read error correction and graph annotation can improve assembly performance of de Bruijn graphs and that this can be seen with the recovery of large (12kbp) events in short read sequences. Moreover, through application to real data we have shown that links can be generated from a wide range of sequencing technologies including data not used to construct the underlying dBG, and that this can be exploited to identify sequences of biological interest. LdBGs can also naturally represent paired-end connectivity information. We have proved that in the error-free setting, Linked de Bruijn Graphs losslessly store the genome sequence, even when constructed from short reads and agnostic of *k*.

Our method is useful for reconstructing complex loci across multiple samples using a common panel of pre-determined haplotypes. Link information derived from a haplotype panel cannot add *k*-mers or edges to the graph that were not observed in the original dataset. Nevertheless, assembly is enhanced in regions where the links are consistent with the graph, and naturally defaults to link-uninformed navigation in regions of discrepancy. Threading a panel of haplotypes from multiple samples through each graph thus identifies only the relevant sections of each donor haplotype.

One shortcoming of long links is the accumulated probability of encountering an error during traversal. If a link takes the wrong branch of an error-induced bubble, cleaning that junction choice trims off all the remaining information about the junction choices made beyond the bubble. This shortcoming results in link coverage dropping off quicker than expected as links get longer, resulting in truncated links. This could be addressed by error-correcting groups of links that start at the same *k*-mer.

We have implemented a very simple read mapping, which trusts all *k*-mers from a read if there is a perfect match in the graph, and which only attempts to fill gaps using linear time graph traversal. Optimal mapping is ultimately NP-hard, but more advanced heuristic methods are available which may perform better than our approach (Limasset *et al.*, 2016). Improvement may be most noticeable for high error rate sequencing data and in low complexity regions of the graph.

Finally, there is scope for reducing memory consumption, given very few *k*-mers actually have links attached (see Supplementary Figure 6) and could be further reduced with better encoding in memory of the junction choice tree held by a *k*-mer (i.e. *L*(*v*)). For example, using a binary encoding of the tree of junction choices, or generative path encoding proposed to compress sequence data (Kingsford and Patro, 2015).

## Acknowledgements

We thank Jerome Kelleher for his many useful comments and edits, Rachel Norris for her pointers and insight into the *K. pneumoniae* dataset, and the other members of the McVean group for useful discussions during the preparation of this manuscript.

